# Conformational variability of cyanobacterial ChlI, the AAA+ motor of magnesium chelatase involved in chlorophyll biosynthesis

**DOI:** 10.1101/2023.04.15.537025

**Authors:** Dmitry Shvarev, Alischa Ira Scholz, Arne Moeller

## Abstract

Magnesium chelatase is a conserved enzyme complex responsible for the first committed step of chlorophyll biosynthesis in photosynthetic organisms, which is the addition of magnesium to the chlorophyll precursor, protoporphyrin IX. The complex is composed of the catalytic subunit ChlH, the bridging subunit ChlD, and the subunit ChlI, which serves as the motor that drives the entire complex. Although the enzyme is well-characterized functionally, high-resolution structures are available only for individual subunits. Hence, the full assembly and the molecular mechanism of the enzyme complex remains unknown. Here, we used cryo-EM, supported by biochemical analysis and mass photometry, to determine structures of the ChlI motor subunit of magnesium chelatase under turnover conditions in the presence of ATP. Our data reveal the molecular details of ChlI oligomerization and conformational dynamics upon ATP binding and hydrolysis. These findings provide new insights into the mechanistic function of ChlI and its implications for the entire magnesium chelatase complex machinery.

## Introduction

Photosynthesis is a vital process in plants, phototrophic bacteria, and cyanobacteria, whereby light energy is absorbed and used to synthesize organic molecules. Essential for photosynthesis are chlorophylls, tetrapyrrole compounds, which harvest the energy of photons. The first committed step in chlorophyll biosynthesis is the insertion of Mg^2+^ into the macrocycle of the chlorophyll precursor protoporphyrin IX ^1,2^, performed by the magnesium chelatase enzyme complex. Magnesium chelatase is fueled by ATP and consists of three subunits: ChlH, ChlD, and ChlI. The motor subunit ChlI belongs to the superfamily of AAA+ (<u>A</u>TPases <u>a</u>ssociated with diverse cellular <u>a</u>ctivities) proteins and, through binding and hydrolyzing of ATP, provides the energy for the thermodynamically unfavorable reaction of Mg^2+^ insertion into protoporphyrin IX ^3,4^. Magnesium chelatase has been extensively characterized functionally ^5,6^, including its activity ^4,7,8^, substrate specificity ^9,10^, and subunit interactions ^7,11–13^. However, how the subunits cooperate and mechanistically contribute to the process of magnesium chelation remains elusive.

Several structures of magnesium chelatase components have recently been reported and provided insights into the molecular organization of the enzyme ^14–16^. The monomeric structure of the AAA+ protein BchI, the ChlI homolog from the photosynthetic bacterium *Rhodobacter capsulatus* (hereafter, *Rhodobacter*), revealed that the protein has an unusual domain arrangement compared to other AAA+ proteins ^14^, placing BchI/ChlI in clade 7 of AAA+ proteins ^17^. As such, the C-terminal domain (small AAA+ subdomain) of BchI is dislocated backward from the nucleotide-binding site, whereas, in other AAA+ proteins, it tends to face upwards. In addition, the protein carries three structured insertions, i1, i2, and i3, located before β-strand β2, within the α-helix α2, and after the α-helix α3. Insertions i2 and i3 can be classified as H2 and PS1 inserts, respectively ^18^. Of note, the insertion i1 is specific for BchI and may play a role in interactions with other components of the magnesium chelatase complex ^14^.

Like other members of the AAA+ superfamily, ChlI/BchI forms oligomers in solution, usually hexamers ^14,15,19^. Recently, the structure of the ChlI hexamer from the cyanobacterium *Synechocystis* sp. PCC 6803 (hereafter, *Synechocystis*) was solved by X-ray crystallography ^15^. In that structure, monomers exhibited well-structured AAA+ large and small subdomains, while insertions i1-i3 were not observed. No nucleotides were bound in the protein, but the ATP-binding pockets at the intermonomer interface showed a typical arrangement for AAA+ proteins.

Moreover, ChlI/BchI forms oligomeric complexes with the other component of magnesium chelatase, ChlD/BchD ^7,11,20^. Previous cryo-EM studies, at intermediate resolution, demonstrated that the BchID subcomplex of magnesium chelatase exhibits a double-ring architecture and undergoes conformational changes upon incubation with different nucleotides ^19,21^. However, no ATPase activity has been detected for ChlD/BchD, despite belonging to the AAA+ superfamily ^7,20^. Apparently, ChlD/BchD only links ChlI/BchI to the porphyrin-binding ChlH/BchH, the third subunit of the complex, leaving the motor function exclusively to ChlI/BchI. Recent biochemical data confirmed that ChlD bridges the ChlI motor to ChlH ^12^. In conjunction, the structural and biochemical data suggest that the ChlI/BchI and ChlD/ChlI rings stack on top of each other and that this assembly hub powers the ChlH/BchH catalytic component of the complex, but how the energy is transferred from ChlI/BchI during this process is unknown.

Here, we used a combination of cryo-EM and functional analysis to investigate the mechanism of the ChlI motor subunit of magnesium chelatase from the cyanobacterium *Nostoc* sp. PCC 7120 (hereafter, *Nostoc*). We examined the oligomerization of ChlI using mass photometry ^22^ and determined the structures of different ChlI oligomeric states with cryo-EM. Our data provide insights into the formation and function of the ChlI hexameric complex, highlighting the ATP-induced conformational changes of the protein. Our results suggest a mechanism whereby the ChlI insertions may facilitate the transfer of ATP binding and hydrolysis energy to the other components of the magnesium chelatase complex.

## Results

### *Purification and biochemical characterization of* Nostoc *ChlI*

We overexpressed the recombinant *Nostoc* ChlI subunit (encoded by the *all0152* gene) in *Escherichia coli* and purified it by affinity and size-exclusion chromatography (Figure 1A). The size-exclusion chromatography profile reveals a single major peak containing pure ChlI protein of approximately 40 kDa, as shown by SDS gel electrophoresis (Figure 1B). We confirmed that ChlI performs ATP hydrolysis in the presence of MgCl_2,_ as expected (Figure 1C).

**Figure 1.**
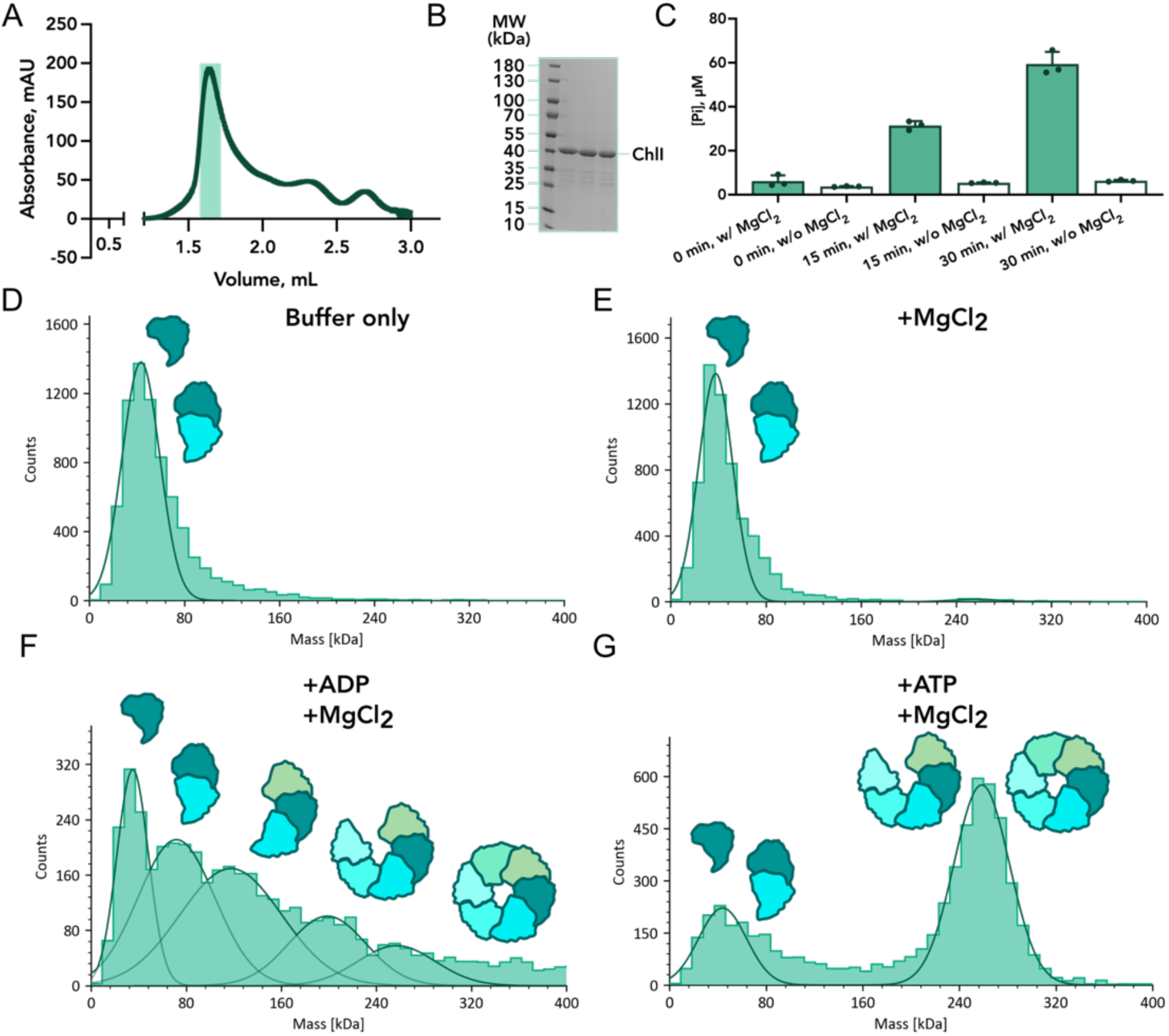
Biochemical and oligomerization analysis of purified ChlI. A) ChlI size-exclusion chromatography (SEC) profile, peak fractions are indicated by a teal rectangle; B) Corresponding SDS electrophoresis gel of SEC peak fractions; C) Phosphate release from ATP hydrolysis by ChlI (12.2 μM) measured using the malachite green assay; D-G) ChlI oligomeric state analyzed in Tris buffer alone (D), with the addition of MgCl_2_ (50 mM) (E), with the addition of ADP (10 mM) and MgCl_2_ (50 mM) (F), or with the addition of ATP (10 mM) and MgCl_2_ (50 mM) (G). For mass photometry, representative results of three independent experiments are shown.

As shown by mass photometry, the protein is mainly monomeric or spontaneously forms small oligomers in solution (Figure 1D, E), which are primarily dimers and are inaccessible by cryo-EM. Therefore, we stimulated the formation of hexamers by incubating the ChlI protein with ATP and MgCl_2_, similar to previous reports on the bacterial ortholog of ChlI, named BchI, and other AAA+ proteins ^14,23,24^. Mass photometry discovered two significant populations of ChlI oligomers that correspond to mono-/dimers and penta-/hexamers in the presence of ATP and MgCl_2_ (Figure 1G). Interestingly, when we added ADP instead of ATP, we observed a wide range of mass distribution in the sample (Figure 1F). We interpret this as a variety of ChlI oligomeric species from monomers to hexamers but with a much smaller fraction of large oligomers.

### Cryo-EM structures of hexameric ChlI

To elucidate the molecular mechanism by which ChlI drives the entire magnesium chelatase complex, we determined the protein structures in the presence of ATP and MgCl_2_ using cryo-EM. For our study, we collected and combined two datasets of approximately 4000 movies each (Figure S1) and processed the data using cryoSPARC ^25^. In support of our biochemical data, we mainly observed the ChlI hexamers in the cryo-EM sample. DTT ^26^ and LMNG were added to combat the otherwise strong preferred particle orientation towards the top/bottom views.

Iterative rounds of 3D classification, performed by multiple-class ab initio reconstructions followed by heterogeneous refinements (Figure S1), revealed two distinct classes, which we named hexamer conformation A and B. These reconstructions achieved global resolutions of 4 Å and 3.8 Å, respectively, with much greater local resolutions in the central parts of the structures (Figure S2). The resulting cryo-EM maps allowed for accurate molecular model building and assignment of nucleotides to their respective densities (Figures 2B, C, 3). Interestingly, each monomer contains a bound nucleotide molecule (ATP or ADP) in both hexameric states. The *Nostoc* ChlI hexameric ring conformations have diameters of approximately 114-117 Å, with the central pore of about 33 Å. Our reconstructions with no symmetry imposed revealed structural differences between the constituent monomers within the ring. Notably, the region of the insertions displayed varying levels of structural organization among the monomers (Figure 2B, C, right panels). This observation differentiates our structures from the previously published structure of *Synechocystis* ChlI ^15^, which did not resolve any structured insertions (Figure S3). Furthermore, in our ChlI hexamers, the monomers are pulled together in the upper region where the insertions are located, making the *Nostoc* ChlI ring more compact in contrast to *Synechocystis*.

**Figure 2.**
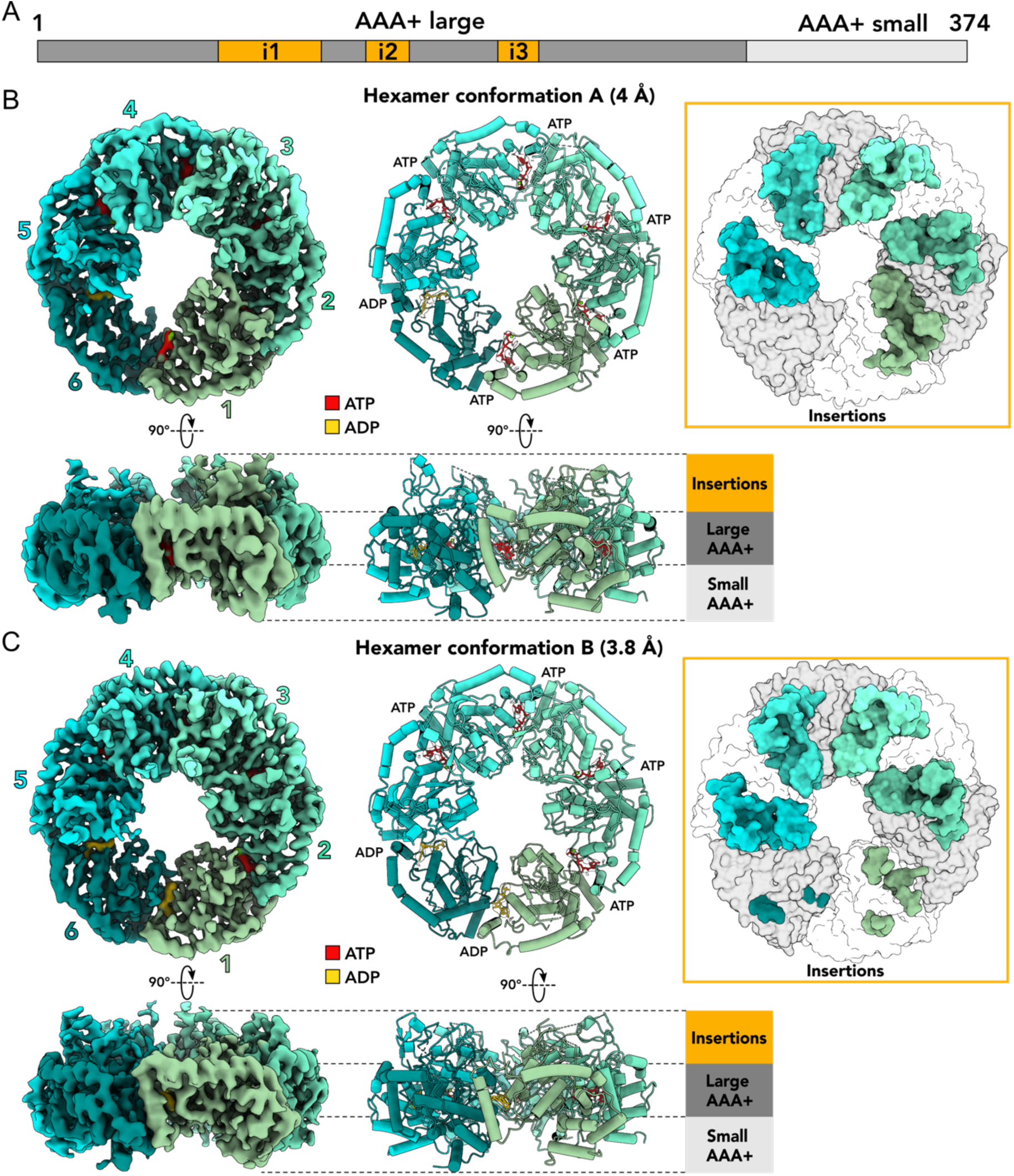
Cryo-EM analysis of ChlI. A) Schematic overview of the ChlI protein sequence with AAA+ subdomains and characteristic insertions i1-i3 indicated; B-C) Cryo-EM maps (left) and corresponding ribbon representations (center) of ChlI hexamer structures in conformation A (panel B) and B (panel C) shown as top and side views. The right panels in (B) and (C) show top views for each hexamer conformation as surface representation with insertions colored by subunit. Densities of modeled nucleotides are shown within the ChlI structure ribbon representations. Large and small AAA+ subdomains and insertions are indicated in the side views of the structures.

**Figure 3.**
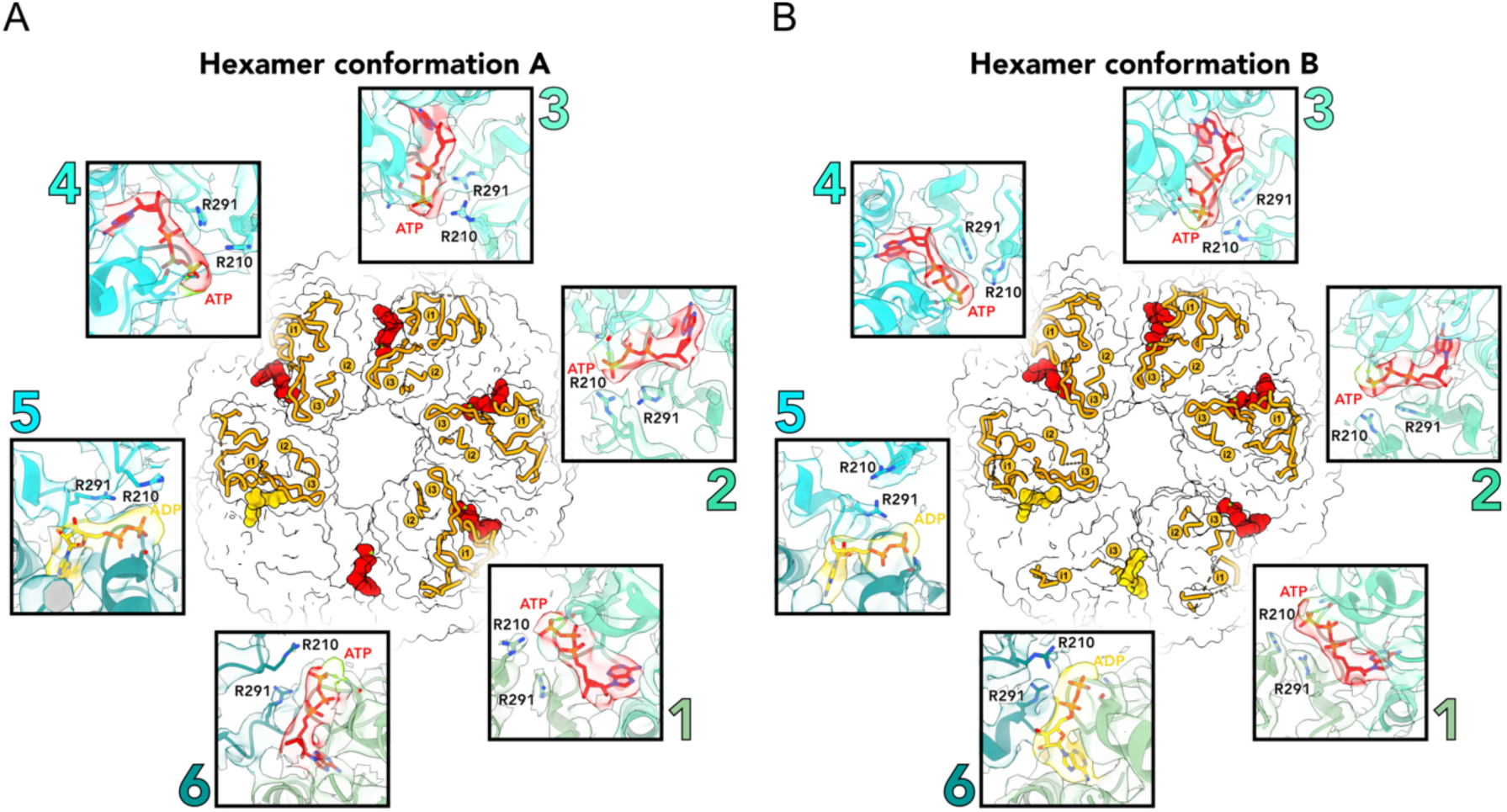
Conformations of ChlI hexamers depend on the bound nucleotides. Molecular surface representations of conformation A (panel A) and conformation B (panel B). Bound ATP (red) and ADP (yellow) molecules are shown as sphere representations inside the hexamers, and structured inserts are shown as ribbons (orange). Insets show close-up views of the nucleotide molecules with surrounding structural elements of ChlI monomers (numbered and colored as in Figure 2); associated cryo-EM densities are zoned around the molecular models and colored accordingly. The Walker A, arginine finger R210, and the sensor 2 R291 residues are indicated.

The core of each *Nostoc* ChlI monomer consists of the large and small AAA+ subdomains resembling a commonly found AAA+ fold ^18,27^. Like in *Rhodobacter* BchI ^14,19^ and *Synechocystis* ChlI ^15^, the *Nostoc* ChlI N-terminal large subdomain consists of six α-helices and five β-strands, and the C-terminal small subdomain consists of four additional α-helices. The small AAA+ subdomain, also called the lid domain, is connected to the large subdomain by α-helices 6 and 7, which can also be considered the presensor 2 insert ^18^, thus assigning ChlI to clade 7 of AAA+ proteins.

ChlI and its close homologs ^14^ contain the structured insertions i1, i2, and i3 (Figures 2, 3) in addition to the standard AAA+ subdomains. Insertion i2 interrupts helix α2 in its first half, and i3 starts immediately after helix α3. Therefore, according to their positions, hairpin insertions i2 and i3 can be attributed to the H2 and PS1 inserts of other AAA+ proteins ^18^. The presence of these insertions further supports the relation of ChlI to clade 7 of AAA+ proteins. On the other hand, the relatively large insert i1 is located immediately after α1 and is more specific for ChlI-like proteins.

### Conformational changes of ChlI upon ATP hydrolysis

We attribute our hexamer structures of ChlI (Figures 2, 3) to its two different states throughout the ATP hydrolysis cycle. Conformation A is characterized by ATP molecules bound in five monomers and ADP in one (Figure 3A), while conformation B has ATP bound in four monomers and ADP in two (Figure 3B). The nucleotide-binding pockets at the interfaces of the monomers demonstrate a standard architecture, including the Walker A and Walker B motifs. Arginine finger R210 and sensor 2 R291 residues from the adjacent monomer additionally coordinate the nucleotide molecule (Figure 3) and contribute to the inter-subunit cooperation within the hexamer, which is typical for AAA+ proteins ^28,29^. In the hexamer structures, the nucleotide state of each monomer is related to the structural organization of the insertions of that monomer, such that when an ATP molecule is bound, the insertions are folded and in an upward position (Figures 2, 3). If ADP is bound, the inserts of this monomer are more flexible or even unfold, as demonstrated by fuzzy or absent corresponding cryo-EM densities (Figure 2). Interestingly, in the observed ADP-bound states, the side chains of the adjacent R210 arginine fingers are positioned somewhat away from the nucleotide (Figure 3A, B inset 5), in line with data from other AAA+ proteins ^29^. In contrast, in most ATP-bound states in our maps, these side chains are oriented towards the nucleotide (Figure 3A, B insets 1-3). This may indicate that these residues also contribute to the coupling of ATP hydrolysis with the reorganization of the inserts in ChlI.

### Pentameric structure of ChlI

Our 3D classifications revealed a class corresponding to a pentameric structure of ChlI (Figure 4), which we attribute to an intermediate state of ChlI complex formation. The map reached an overall resolution of 4.9 Å and differs significantly from the hexamer maps (Figures 2, 3). The ChlI pentamer shows a spring-washer-like helical organization with an opening of approximately 45 Å between the first and last monomer in the complex, which would be sufficient to accommodate an additional monomer of ChlI. The resolution obtained for this structure allowed for the fitting of α-helices and β-strands of ChlI monomers but limited the determination of bound nucleotide molecules. The three central monomers of the pentamer contain partially folded insertions (Figure 4B) and generally show higher local resolution (Figure S2C). In contrast, the two peripheral monomers are less resolved, possibly due to the increased flexibility of the ChlI pentamer at its ends.

**Figure 4.**
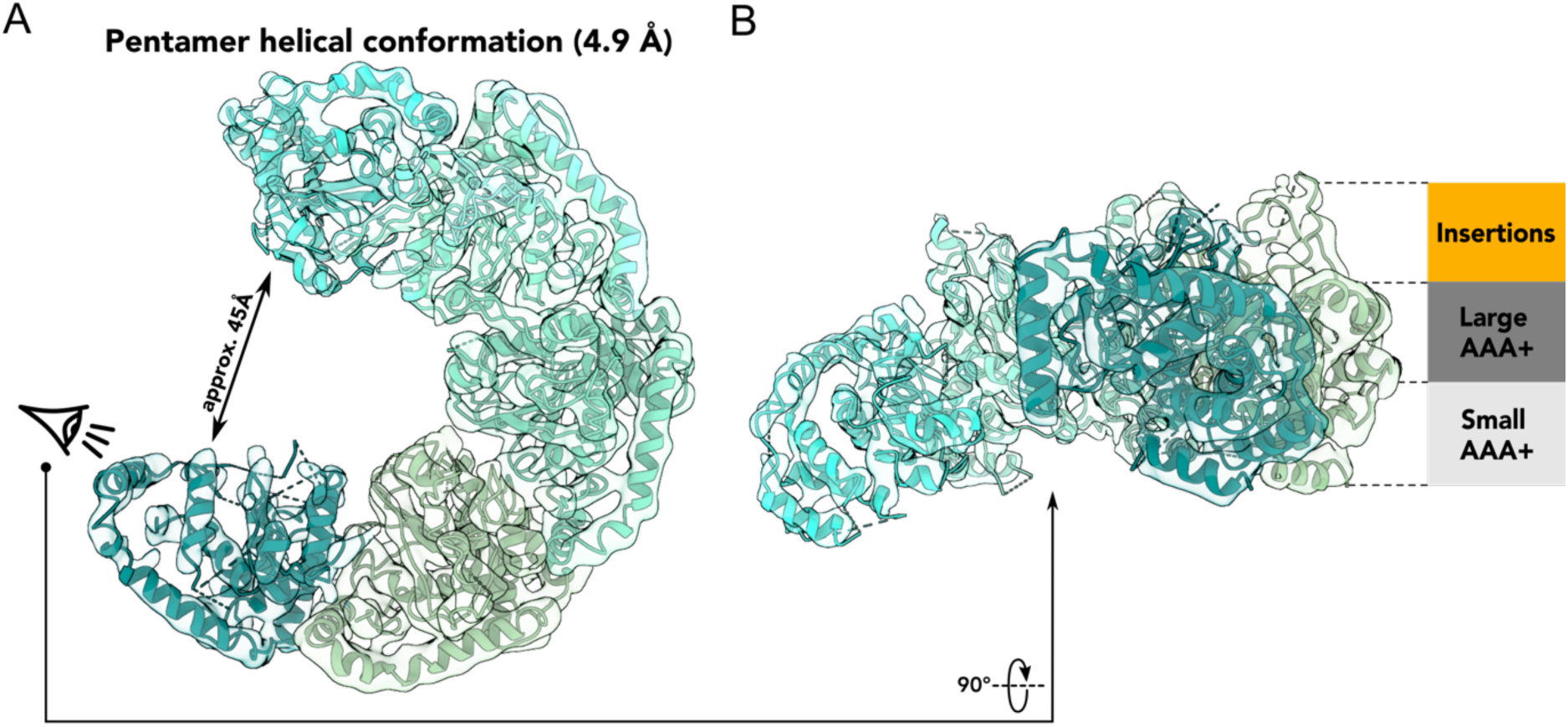
Pentameric spring-washer-like structure of ChlI. A) Top view of the cryo-EM density (transparent) and corresponding molecular model in ribbon representation colored by subunit as in Figure 2; B) Helical ChlI pentamer viewed from the side; positions of the insertions (orange) and AAA+ subdomains (gray) are indicated.

## Discussion

In this study, we analyzed the ChlI motor subunit of the magnesium chelatase from the filamentous nitrogen-fixing cyanobacterium *Nostoc* sp. PCC 7120 ^30,31^. Using biochemical methods and cryo-EM, we discovered two asymmetric hexameric states and one pentameric state of ChlI in the presence of ATP (Figures 1-4). Mass photometry showed that ATP induces the formation of large ChlI oligomers such as hexamers and pentamers (Figure 1G), as previously reported for ChlI homologs ^19,24^. Interestingly, adding ADP also stimulates oligomer formation by ChlI, but it is insufficient for stable hexamer formation (Figure 1F).

The ChlI monomers in our structures show similarities to other AAA+ proteins and previously characterized ChlI-like proteins ^18,27–29^, including BchI from *Rhodobacter* ^14^ and ChlI from *Synechocystis* ^15^ (Figure S3). However, our hexameric ChlI structures exhibit unique variations in the architecture of the insertion domains, dependent on the nucleotide state of the monomers, which is not observed previously in ChlI close homologs. The insertions are folded upwards when a monomer is ATP-bound and are more flexible or disordered when ADP is bound (Figures 2, 3). It has been shown for AAA+ proteins dynein and CbbQO-type Rubisco activase that their H2 and PSI inserts have an essential mechanistic function since they bind to partner proteins and couple this with ATP hydrolysis ^32–34^. Similarly, we suggest that the alternate appearance of insertions i1-i3 in ChlI may be essential for interactions with other subunits and thus for the mechanism of magnesium chelatase.

Based on our data, we propose a sequence of events that ChlI undergoes in the presence of ATP (Figure 5). First, ChlI monomers interact to form small oligomers upon ATP binding or spontaneously (Figure 1D,E). The oligomers then grow and form helical spring-washer-like pentameric structures, potentially representing a resting state of the enzyme (Figures 1G, 4), similar to the Lon protease’s open helical state ^35^ or the AAA+ protein RavA, which creates a mix of open spiral and closed planar ring state populations ^36^. Upon binding another monomer to the pentamer, ChlI undergoes rigid-body rearrangements to form a nearly planar hexameric ring. This addition of the sixth monomer to the complex would then initiate ATP hydrolysis, as proposed for Vps4 ^37^.

**Figure 5.**
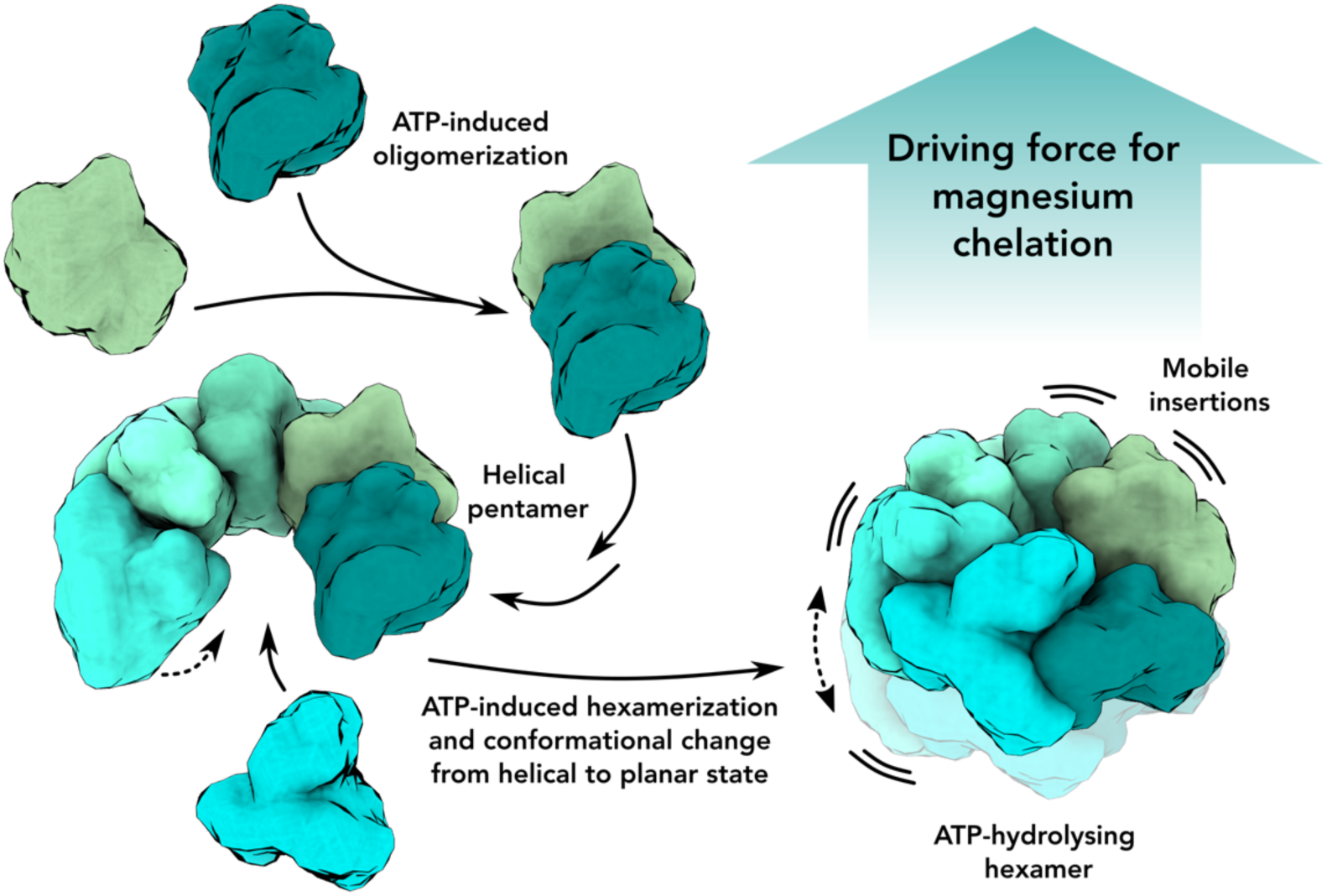
Model of ChlI oligomerization and mechanism. In the presence of ATP, ChlI monomers begin to oligomerize and eventually form a stable spring-washer-like helical pentameric structure. Once the sixth monomer and the sixth molecule of ATP bind to the pentamer, the hexamer ring is formed, and ATP hydrolysis is initiated. The ATP hydrolysis cycle, which proceeds from one monomer to the other, increases the flexibility of the insertions and forces the ChlI ring from a helical to a planar conformation. The conformational changes in the insertions and the entire ChlI ring probably provide the driving force for the functioning of the whole magnesium chelatase complex.

According to our model, ATP hydrolysis occurs in one monomer at a time, triggering rearrangements involving residues near the nucleotide-binding pocket. This induces reorganization of the monomer’s insertions and flattening of the entire hexamer (Figures 2, 3), which may contribute to ATP hydrolysis in the adjacent monomer, similar to the mechanism of peptide-translocating AAA+ proteins ^27^. We speculate that after all the monomers have converted ATP to ADP, the ring disassembles for nucleotide exchange. Nevertheless, additional structural data is needed to understand ATP hydrolysis in ChlI.

The role of the insertion domains in ChlI-ChlD interactions is still unclear and also requires further investigation. It has been proposed that ChlI forms a two-tiered ring complex after binding to ChlD ^14,15^, similar to the type II AAA+ proteins ^27^. Moreover, the corresponding low-resolution structure of the BchI-BchD double-ring complex has been determined ^19^. However, biochemical data show that arginine finger and sensor 2 residues and the entire AAA+ module of ChlD are essential for binding to ChlI in *Synechocystis* ^11^. This may indicate that ChlD and ChlI form a mixed hetero-hexameric one-tiered ring complex. We have performed AlphaFold ^38,39^ modeling of complex formation between the monomers of ChlI and ChlD from *Nostoc* and show in our prediction that the ChlD monomer has an almost identical interface with ChlI as the interface between ChlI monomers in our cryo-EM structures (Figure S4). This observation may support the idea that ChlI and ChlD form a mixed ring within magnesium chelatase.

In conclusion, our structures provide insights into the oligomerization and ATP hydrolysis mechanism of the clade 7 AAA+ protein ChlI. According to our mechanistic model (Figure 5), in the context of the entire magnesium chelatase complex, ATP hydrolysis by ChlI would induce conformational changes in the insertions of its monomers, leading to structural reorganization of ChlD protomer(s) bound to ChlI ^11,19^. This reorganization, consequently, would transfer the energy of ATP hydrolysis to ChlH, the catalytic subunit of magnesium chelatase ^16^ interacting with ChlD ^12^.

## Materials and methods

### Protein overproduction and purification

The gene *all0151* was amplified from the genomic DNA of *Nostoc* sp. PCC 7120 using primers purchased from Biolegio with the sequences 5’–ATT TTG TTT AAC TTT AAG AAG GAG ATA TAC ATG CAT CAT CAT CAT CAT CAC ACT CCA ACA GCT CAA ACC ACG G–3’ and 5’–GTG ATG ATG ATG ATG ATG GCT GCT GCC CAT GTT ACC GGA CAC CTG TCT TAA TTT G–3’, was cloned using Gibson assembly ^40^ into plasmid pET15b (Novagen/Merck), which was digested with NcoI. The N-terminal His-tag was introduced into the sequence. The protein was overproduced in *E. coli* LOBSTR cells ^41^. The cells were grown in Luria broth (LB) medium at 37°C with continuous shaking. Protein synthesis was induced by adding 0.1 mM isopropyl-β-d-thiogalactoside (IPTG) at an OD^600^ of 0.6 and incubating the cultures overnight at 18°C with shaking. After induction, cells were harvested by centrifugation at 5,000 *g* for 20 minutes at 4°C. The pellet was resuspended in Tris buffer (20 mM Tris, 200 mM NaCl, pH 7.5) containing 1 mM phenylmethylsulfonyl fluoride (PMSF) and 1 mg/ml lysozyme and incubated at 4°C for 1-2 hours with shaking. Then, 1 mg/ml of DNAse I and 5 mM MgCl_2_ were added, and the suspension was incubated for another 1-2 hours. The solutions were centrifuged at 17,000 *g* for 30 min at 4°C. The supernatant containing extracted soluble proteins was used to purify recombinant ChlI by affinity chromatography using a gravity flow column with nickel-nitriloacetic acid resin (Thermo Scientific). The column was equilibrated with 10 column volumes of Tris buffer (see above) containing 10 mM imidazole. The lysate was applied and incubated on the column for 2 hours, followed by washing with Tris buffer containing 20 mM imidazole, and ChlI was eluted with Tris buffer containing 250 mM imidazole. The concentrated eluate was applied to a Superdex 200 Increase 15/150 GL column (Cytiva) for size exclusion chromatography (SEC) and eluted in 50 μL fractions using an ÄKTAgo purification system (Cytiva). Peak fractions were used for further analysis.

### ATP hydrolysis assays

The ATP hydrolysis activity of ChlI was assessed by quantification of free orthophosphate using the malachite green phosphate assay kit (Sigma-Aldrich) according to the manufacturer’s protocol. Briefly, ChlI at 0.5 mg/ml (12.2 μM) was incubated in Tris buffer (see above) with 10 mM of ATP and with or without the addition of 50 mM MgCl_2_ for different time intervals at 37°C. After incubation, the samples were diluted until the ATP concentration reached 0.25 mM and immediately frozen in liquid nitrogen to stop the reaction. The samples were then thawed, transferred to a 96-well plate, and treated with the malachite green reagent. The plate was incubated for additional 30 minutes at room temperature for color development, and absorbance at 620 nm was measured using a SpectraMax M3 plate reader (Molecular Devices). Phosphate concentrations in the samples were determined from a previously generated standard curve.

### Mass photometry

Mass photometry ^22^ experiments were performed using a Refeyn TwoMP instrument (Refeyn Ltd.). Data were obtained using AcquireMP software and analyzed using DiscoverMP (both Refeyn Ltd.). Glass coverslips were used for sample analysis. Perforated silicone gaskets were placed on the coverslips to form wells for each sample to be measured. ChlI samples were incubated at a concentration of 0.5 mg/ml in Tris buffer (see above) with various additives (Figure 1D-G). Samples were further diluted 1:5 immediately prior to analysis, and 2 μL of this solution was added to 18 μL of Tris buffer placed in the gasket for the measurement.

### Cryo-EM sample preparation and data acquisition

Prior to cryo-EM, ChlI samples were examined by negative-stain electron microscopy, using 2% (w/v) uranyl formate solution as previously described ^42^. Negative-stain micrographs were recorded manually on a JEM-2100Plus transmission electron microscope (JEOL) operated at 200 kV and equipped with a XAROSA CMOS 20-megapixel camera (EMSIS) at a nominal magnification of 30,000 (3.12 Å per pixel).

For cryo-EM, 3 μL of 0.5 mg/ml of freshly purified ChlI with the addition of 10 mM ATP, 50 mM MgCl_2_, 0.05% lauryl maltose neopentyl glycol (LMNG), and 1 mM dithiothreitol (DTT) were applied to glow-discharged C-Flat grids (R1.2/1.3 3Cu-50) (EMS) and immediately plunge frozen in liquid ethane using a Vitrobot Mark IV (Thermo Fisher Scientific) with the environmental chamber set at 100% humidity and 4°C.

Two data sets of 4191 and 3495 movies were recorded automatically with EPU (Thermo Fisher Scientific), using a Glacios cryo-transmission electron microscope (Thermo Fisher Scientific) operated at 200 kV and equipped with a Selectris energy filter and a Falcon 4 detector (both Thermo Fisher Scientific). Data were recorded in Electron Event Representation (EER) mode at a nominal magnification of 130,000 (0.924 Å per pixel) in the defocus range of –0.8 to –1.8 μm with an exposure time of 7.50 s resulting in a total electron dose of approximately 50 e^−^ Å^−2^.

### Cryo-EM image processing

All cryo-EM data were preprocessed in cryoSPARC Live, and further processing was performed in cryoSPARC v3 and v4 ^25^ (Figure S1). For all collected movies, motion correction (EER upsampling factor 2, EER number of fractions 40) and contrast transfer function (CTF) estimation were performed using cryoSPARC Live implementations. Both collected datasets were preprocessed in a similar manner and combined at the final image processing steps (Figure S1). For the dataset of 4191 micrographs, particles were selected by the template picker implemented in CryoSPARC Live, using well-defined 2D classes obtained from previous ChlI datasets as templates. Picked particles were extracted in a box size of 288 pixels, and Fourier cropped to 144 pixels (resulting in a pixel size of 1.848 Å per pixel) and subjected to 2D classification, which produced 2D classes that were used as templates for another round of template particle picking in CryoSPARC followed by particle extraction using the same box size of 288 pixels (Fourier cropped to 144 pixels) and several rounds of 2D classification to eliminate bad picks. In addition, particles were picked using the Topaz ^43^ wrapper in CryoSPARC and also extracted using a box size of 288 pixels (Fourier cropped to 144 pixels). All picked particles were then combined, and duplicates were removed, resulting in a stack of 581.4K particles. These particles were then subjected to two rounds of 2D classification. The 3495-movie dataset was processed similarly, and all particle pickers applied there resulted in a stack of 375.2K particles after duplicate removal (Figure S1). These particles were also subjected to two rounds of 2D classification.

Particles from the two datasets were then combined and used for four rounds of ab-initio reconstructions with multiple classes followed by heterogeneous refinements. The two best hexamer classes (80.5K and 58.3K particles) and the class corresponding to the pentamer (24.7K particles) from the last round of heterogeneous refinement were further refined by non-uniform refinement ^44^ using particles re-extracted with the full box size (288 pixels, 0.924 Å per pixel), resulting in reconstructions with resolutions of 3.9 Å and 4.1 Å for the hexamers and 5,6 Å for the pentamer (Gold Standard Fourier Shell Correlation (GSFSC) value of 0.143). Consequently, the reconstructions were further improved using the non-uniform refinement with CTF refinement on the fly resulting in consensus maps with resolutions of 3.8 Å and 4 Å for the hexamers and 4.9 Å for the pentamer (GSFSC = 0.143) (Figure S1).

All maps were subjected to unsupervised B-factor sharpening within cryoSPARC. No symmetry was applied during processing. The quality of the consensus maps is shown in Figure S2. All GSFSC curves were generated within cryoSPARC. The local resolutions of the consensus maps (Figure S2) were estimated in cryoSPARC and analyzed in UCSF ChimeraX ^45^. Dataset statistics are provided in Table S1.

### Model building and refinement

For each of the three structures, the same approach has been used to build the model. The AlphaFold structure prediction of the ChlI monomer (AlphaFold accession: AF-P58571-F1; Uniprot: P58571) was manually fitted to each of three maps of ChlI using the “Fit in Map” tool in ChimeraX ^45^ and used as a starting model. The N-terminal (residues 1-13) and C-terminal (residues 354-374) fragments of ChlI without well-resolved densities were removed. The models of the ChlI monomer were then manually adjusted and refined in Coot ^46^. The models of the other monomers of ChlI were generated based on the built structure of the first monomer and fitted to the ChlI cryo-EM maps using UCSF ChimeraX. Models of individual monomers were then combined into a single structure and manually inspected and refined in Coot, and models of respective nucleotides and magnesium ions were fitted inside the corresponding densities. The fragments of ChlI monomers with missing corresponding densities in the cryo-EM maps were manually deleted in Coot. Subsequently, iterative rounds of real space refinement ^47^ of the models against the corresponding ChlI maps in PHENIX ^48^ were performed, followed by manual adjustments in Coot. The model of the ChlI pentamer was truncated to poly-alanine using the PDB Tools job in PHENIX. Model validation was done using MolProbity ^49^ in PHENIX. Models and maps were visualized, and figures were prepared in UCSF ChimeraX and Inkscape. Model refinement and validation statistics are provided in Table S1.

## Acknowledgments

We thank Marvin Wortmann and Rene Leffers for their early contributions, Dovile Januliene for help with cryo-EM, Kristian Parey for help with model building, and Kilian Schnelle for computer support. We also thank Iris Maldener and Karl Forchhammer for their advice.

## Funding

This work was funded by Deutsche Forschungsgemeinschaft CRC944, Deutsche Forschungsgemeinschaft CRC1557, Deutsche Forschungsgemeinschaft INST 190/196-1 FUGG, and Bundesministerium für Bildung und Forschung JPND-DLR 01ED2010.

## Author contributions

Investigation: DS, AIS. Formal analysis: DS, AIS. Visualization: DS. Conceptualization: DS, AM. Funding acquisition: AM. Writing—original draft: DS. Writing—review and editing: DS, AIS, AM.

## Conflict of interests

The authors declare no conflict of interest.

## Data availability

The cryo-EM density maps and corresponding atomic models reported in this study have been deposited in the Electron Microscopy Data Bank and Protein Data Bank with the accession codes EMD-xxx and PDB-xxx, respectively.

## Notes

### Competing Interest Statement

The authors have declared no competing interest.

## References

1. Masuda, T. & Fujita, Y. Regulation and evolution of chlorophyll metabolism. Photochem. Photobiol. Sci. 7, 1131–1149 (2008).

2. Bryant, D. A., Hunter, C. N. & Warren, M. J. Biosynthesis of the modified tetrapyrroles— the pigments of life. J. Biol. Chem. 295, 6888–6925 (2020).

3. Reid, J. D. & Hunter, C. N. Magnesium-dependent ATPase activity and cooperativity of magnesium chelatase from Synechocystis sp. PCC6803. J. Biol. Chem. 279, 26893–26899 (2004).

4. Adams, N. B. P., Brindley, A. A., Hunter, C. N. & Reid, J. D. The catalytic power of magnesium chelatase: a benchmark for the AAA+ ATPases. FEBS Lett. 590, 1687–1693 (2016).

5. Gibson, L. C. D., Willows, R. D., Kannangara, C. G., Von Wettstein, D. & Hunter, C. N. Magnesium-protoporphyrin chelatase of Rhodobacter sphaeroides: Reconstitution of activity by combining the products of the bchH, -I, and -D genes expressed in Escherichia coli. Proc. Natl. Acad. Sci. U. S. A. 92, 1941–1944 (1995).

6. Jensen, P. E., Gibson, L. C. D., Henningsen, K. W. & Hunter, C. N. Expression of the chlI, chlD, and chlH genes from the cyanobacterium Synechocystis PCC6803 in Escherichia coli and demonstration that the three cognate proteins are required for magnesium-protoporphyrin chelatase activity. J. Biol. Chem. 271, 16662–16667 (1996).

7. Jensen, P. E., Gibson, L. C. D. & Hunter, C. N. ATPase activity associated with the magnesium-protoporphyrin IX chelatase enzyme of Synechocystis PCC6803: Evidence for ATP hydrolysis during Mg2+ insertion, and the MgATP-dependent interaction of the ChlI and ChlD subunits. Biochem. J. 339, 127–134 (1999).

8. Reid, J. D., Siebert, C. A., Bullough, P. A. & Hunter, C. N. The ATPase activity of the ChlI subunit of magnesium chelatase and formation of a heptameric AAA+ ring. Biochemistry 42, 6912–6920 (2003).

9. Karger, G. A., Reid, J. D. & Hunter, C. N. Characterization of the binding of deuteroporphyrin IX to the magnesium chelatase H subunit and spectroscopic properties of the complex. Biochemistry 40, 9291–9299 (2001).

10. Sirijovski, N. et al. Substrate-binding model of the chlorophyll biosynthetic magnesium chelatase BchH subunit. J. Biol. Chem. 283, 11652–11660 (2008).

11. Adams, N. B. P., Vasilev, C., Brindley, A. A. & Hunter, C. N. Nanomechanical and Thermophoretic Analyses of the Nucleotide-Dependent Interactions between the AAA+ Subunits of Magnesium Chelatase. J. Am. Chem. Soc. 138, 6591–6597 (2016).

12. Farmer, D. A. et al. The ChlD subunit links the motor and porphyrin binding subunits of magnesium chelatase. Biochem. J. 476, 1875–1887 (2019).

13. Adams, N. B. P. & Reid, J. D. The allosteric role of the AAA+ domain of ChlD protein from the magnesium chelatase of synechocystis species PCC 6803. J. Biol. Chem. 288, 28727–28732 (2013).

14. Fodje, M. N. et al. Interplay between an AAA module and an integrin I domain may regulate the function of magnesium chelatase. J. Mol. Biol. 311, 111–122 (2001).

15. Gao, Y. S., Wang, Y. L., Wang, X. & Liu, L. Hexameric structure of the ATPase motor subunit of magnesium chelatase in chlorophyll biosynthesis. Protein Sci. 29, 1040–1046 (2020).

16. Adams, N. B. P. et al. The active site of magnesium chelatase. Nat. Plants (2020). doi:10.1038/s41477-020-00806-9

17. Erzberger, J. P. & Berger, J. M. EVOLUTIONARY RELATIONSHIPS AND STRUCTURAL MECHANISMS OF AAA+ PROTEINS. https://doi.org/10.1146/annurev.biophys.35.040405.101933 35, p93–114 (2006).

18. Jessop, M., Felix, J. & Gutsche, I. AAA+ ATPases: structural insertions under the magnifying glass. Curr. Opin. Struct. Biol. 66, 119–128 (2021).

19. Lundqvist, J. et al. ATP-Induced Conformational Dynamics in the AAA+ Motor Unit of Magnesium Chelatase. Structure 18, 354–365 (2010).

20. Gibson, L. C. D., Jensen, P. E. & Hunter, C. N. Magnesium chelatase from Rhodobacter sphaeroides: initial characterization of the enzyme using purified subunits and evidence for a BchI-BchD complex. Biochem. J. 337, 243 (1999).

21. Elmlund, H. et al. A New Cryo-EM Single-Particle Ab Initio Reconstruction Method Visualizes Secondary Structure Elements in an ATP-Fueled AAA+ Motor. J. Mol. Biol. 375, 934–947 (2008).

22. Young, G. et al. Quantitative mass imaging of single biological macromolecules. Science (80-.). 360, 423–427 (2018).

23. Lupas, A. N. & Martin, J. AAA proteins. Curr. Opin. Struct. Biol. 12, 746–753 (2002).

24. Willows, R. D., Hansson, A., Birch, D., Al-Karadaghi, S. & Hansson, M. EM single particle analysis of the ATP-dependent BchI complex of magnesium chelatase: An AAA+ hexamer. J. Struct. Biol. 146, 227–233 (2004).

25. Punjani, A., Rubinstein, J. L., Fleet, D. J. & Brubaker, M. A. CryoSPARC: Algorithms for rapid unsupervised cryo-EM structure determination. Nat. Methods 14, 290–296 (2017).

26. Klebl, D. P., Wang, Y., Sobott, F., Thompson, R. F. & Muench, S. P. It started with a Cys: Spontaneous cysteine modification during cryo-EM grid preparation. Front. Mol. Biosci. 9, 767 (2022).

27. Puchades, C., Sandate, C. R. & Lander, G. C. The molecular principles governing the activity and functional diversity of AAA+ proteins. Nat. Rev. Mol. Cell Biol. 2019 211 21, 43–58 (2019).

28. Hanson, P. I. & Whiteheart, S. W. AAA+ proteins: Have engine, will work. Nat. Rev. Mol. Cell Biol. 6, 519–529 (2005).

29. Wendler, P., Ciniawsky, S., Kock, M. & Kube, S. Structure and function of the AAA+ nucleotide binding pocket. Biochim. Biophys. Acta - Mol. Cell Res. 1823, 2–14 (2012).

30. Rippka, R., Deruelles, J., Waterbury, J. B., Herdman, M. & Stanier, R. Y. Generic Assignments, Strain Histories and Properties of Pure Cultures of Cyanobacteria. Microbiology 111, 1–61 (1979).

31. Maldener, I., Summers, M. L. & Sukenik., A. Cellular differentiation in filamentous cyanobacteria. in The Cell Biology of Cyanobacteria (eds. Flores, E. & Herrero, A.) 263–291 (Caister Academic Press, 2014).

32. Tsai, Y. C. C. et al. Insights into the mechanism and regulation of the CbbQO-type Rubisco activase, a MoxR AAA+ ATPase. Proc. Natl. Acad. Sci. U. S. A. 117, 381–387 (2020).

33. Kon, T. et al. The 2.8 Å crystal structure of the dynein motor domain. Nat. 2012 4847394 484, 345–350 (2012).

34. Bhabha, G. et al. Allosteric Communication in the Dynein Motor Domain. Cell 159, 857–868 (2014).

35. Shin, M. et al. Structural basis for distinct operational modes and protease activation in aaa+ protease lon. Sci. Adv. 6, (2020).

36. Jessop, M. et al. Structural insights into ATP hydrolysis by the MoxR ATPase RavA and the LdcI-RavA cage-like complex. Commun. Biol. 3, 1–14 (2020).

37. Han, H. et al. Structure of Vps4 with circular peptides and implications for translocation of two polypeptide chains by AAA+ ATPases. Elife 8, 1–20 (2019).

38. Jumper, J. et al. Highly accurate protein structure prediction. Nature (2021). doi:10.1038/s41586-021-03819-2

39. Evans, R. et al. Protein complex prediction with AlphaFold-Multimer. bioRxiv 2021.10.04.463034 (2021). doi:10.1101/2021.10.04.463034

40. Gibson, D. G. et al. Enzymatic assembly of DNA molecules up to several hundred kilobases. Nat. Methods 6, 343–345 (2009).

41. Andersen, K. R., Leksa, N. C. & Schwartz, T. U. Optimized E. coli expression strain LOBSTR eliminates common contaminants from His-tag purification. Proteins Struct. Funct. Bioinforma. 81, 1857–1861 (2013).

42. Januliene, D. & Moeller, A. Single-Particle Cryo-EM of Membrane Proteins. in Structure and Function of Membrane Proteins, Methods in Molecular Biology 2302, 153–178 (2021).

43. Bepler, T. et al. Positive-unlabeled convolutional neural networks for particle picking in cryo-electron micrographs. Nat. Methods 2019 1611 16, 1153–1160 (2019).

44. Punjani, A., Zhang, H. & Fleet, D. J. Non-uniform refinement: Adaptive regularization improves single particle cryo-EM reconstruction. Nat. Methods 17, 1214–1221 (2020).

45. Pettersen, E. F. et al. UCSF ChimeraX: Structure visualization for researchers, educators, and developers. Protein Sci. 30, 70–82 (2021).

46. Emsley, P., Lohkamp, B., Scott, W. G. & Cowtan, K. Features and development of Coot. urn:issn:0907-4449 66, 486–501 (2010).

47. Afonine, P. V. et al. Real-space refinement in PHENIX for cryo-EM and crystallography. urn:issn:2059-7983 74, 531–544 (2018).

48. Liebschner, D. et al. Macromolecular structure determination using X-rays, neutrons and electrons: recent developments in Phenix. urn:issn:2059-7983 75, 861–877 (2019).

49. Williams, C. J. et al. MolProbity: More and better reference data for improved all-atom structure validation. Protein Sci. 27, 293–315 (2018).

