## Supplemental Material for "Conformational variability of cyanobacterial ChlI, the AAA+ motor of magnesium chelatase involved in chlorophyll biosynthesis"

### Supplementary Data

#### Table S1. Cryo-EM data collection, refinement and validation statistics

| | Hexamer conformation A (EMDB-xxxx) (PDB xxxx) | Hexamer conformation B (EMDB-xxxx) (PDB xxxx) | Pentamer conformation (EMDB-xxxx) (PDB xxxx) |
|---|---|---|---|
| **Data collection and processing** | | | |
| Magnification | 130,000 | 130,000 | 130,000 |
| Voltage (kV) | 200 | 200 | 200 |
| Electron exposure (e–/$Å^2$) | 50 | 50 | 50 |
| Defocus range (μm) | -0.8 to -2.8 | -0.8 to -2.8 | -0.8 to -2.8 |
| Pixel size (Å) | 0.924 | 0.924 | 0.924 |
| Symmetry imposed | C1 | C1 | C1 |
| Initial particle images (no.) | 4,940,012 | 4,940,012 | 4,940,012 |
| Final particle images (no.) | 58,137 | 80,344 | 24,589 |
| Map resolution (Å) | 4.0 | 3.8 | 4.9 |
| FSC threshold | 0.143 | 0.143 | 0.143 |
| **Refinement** | | | |
| Initial model used | AlphaFold | AlphaFold | AlphaFold |
| Model resolution range (Å) | 3.0-6.0 | 3.0-6.0 | 4.0-7.0 |
| FSC threshold | 0.143 | 0.143 | 0.143 |
| Map sharpening *B* factor ($Å^2$) | -103.3 | -105.8 | -189.5 |
| Model composition | | | |
| Non-hydrogen atoms | 14529 | 14536 | 6800 |
| Protein residues | 1829 | 1833 | 1378 |
| Ligands | MG (5), ATP (5), ADP (1) | MG (4), ATP (4), ADP (2) | - |
| R.m.s. deviations | | | |
| Bond lengths (Å) | 0.004 | 0.004 | 0.006 |
| Bond angles (°) | 0.747 | 0.729 | 0.943 |
| Validation | | | |
| MolProbity score | 2.31 | 2.00 | 1.91 |
| Clashscore | 16.26 | 9.67 | 5.91 |
| Poor rotamers (%) | 0.69 | 0.19 | 0 |
| Ramachandran plot | | | |
| Favored (%) | 87.95 | 91.95 | 88.66 |
| Allowed (%) | 12.05 | 7.99 | 11.26 |
| Outliers (%) | 0 | 0.06 | 0.08 |

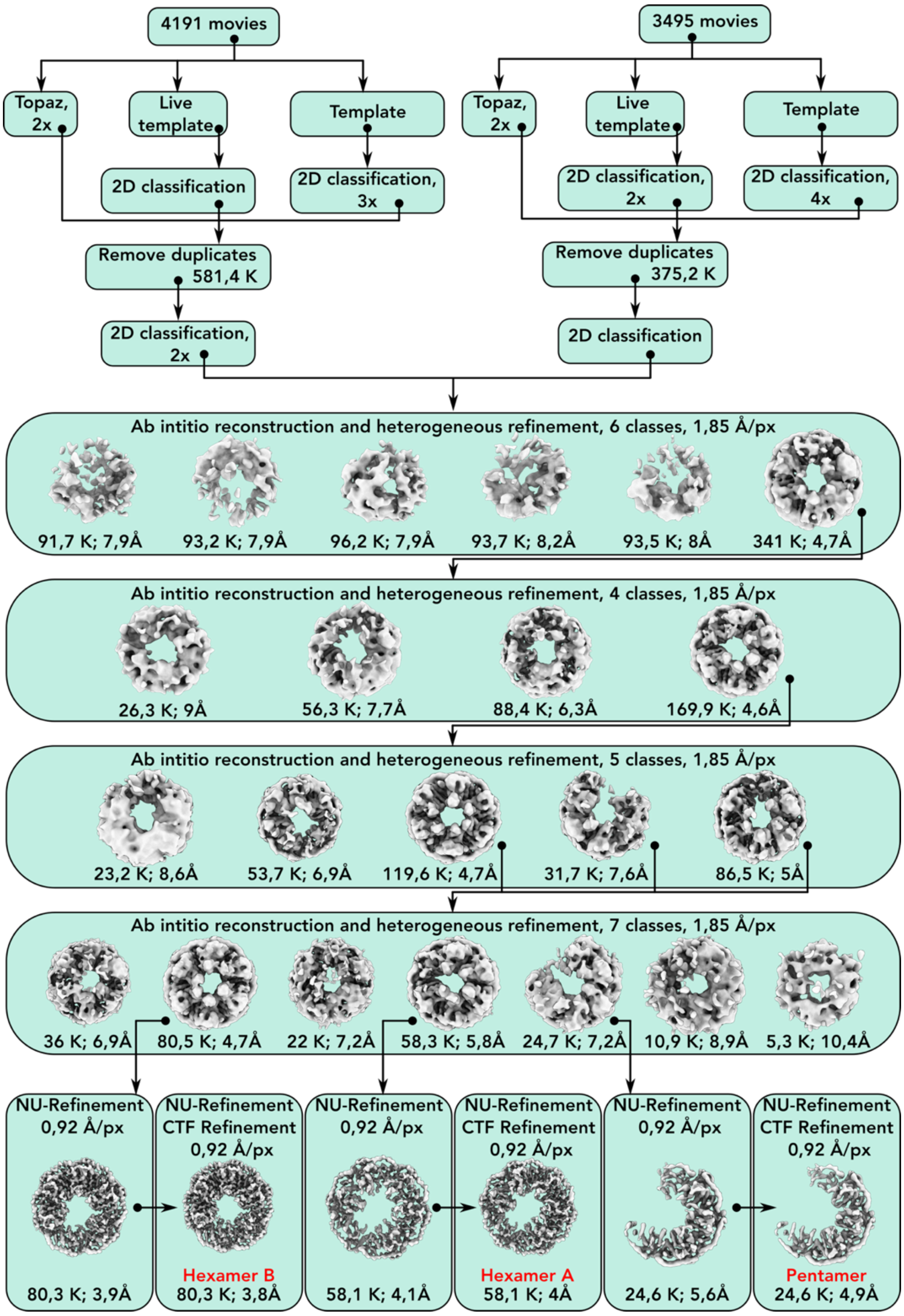

**Figure S1.** Scheme of cryo-EM image processing. A detailed description of the data analysis can be found in the Materials and Methods section. The entire pipeline was performed in CryoSPARC v.3 and v.4. Briefly, two datasets of approximately 4000 movies were collected. After preprocessing in cryoSPARC Live, particles were picked using the template and Topaz pickers. For the template picker, 2D classes from our trial attempts of ChlI cryo-EM analysis were used as templates. After particle picking, the particles were 2D classified, and duplicate particles were removed. Next, additional 2D classification was performed, and the retained particles were combined and subjected to several rounds of ab initio reconstruction and heterogeneous refinement with multiple classes. Particles from the two best ring classes and the pentamer class from the last round of heterogeneous refinement were separately re-extracted using a full box (0.92 Å/pixel) and subjected to a round of NU-refinement for each class. Another round of NU-refinement with CTF refinement was used to further improve the maps. The final maps achieved resolutions of 3.8 Å, 4 Å (for hexamers) and 4.9 Å (for pentamer). No symmetry was applied at any stage of the processing pipeline.

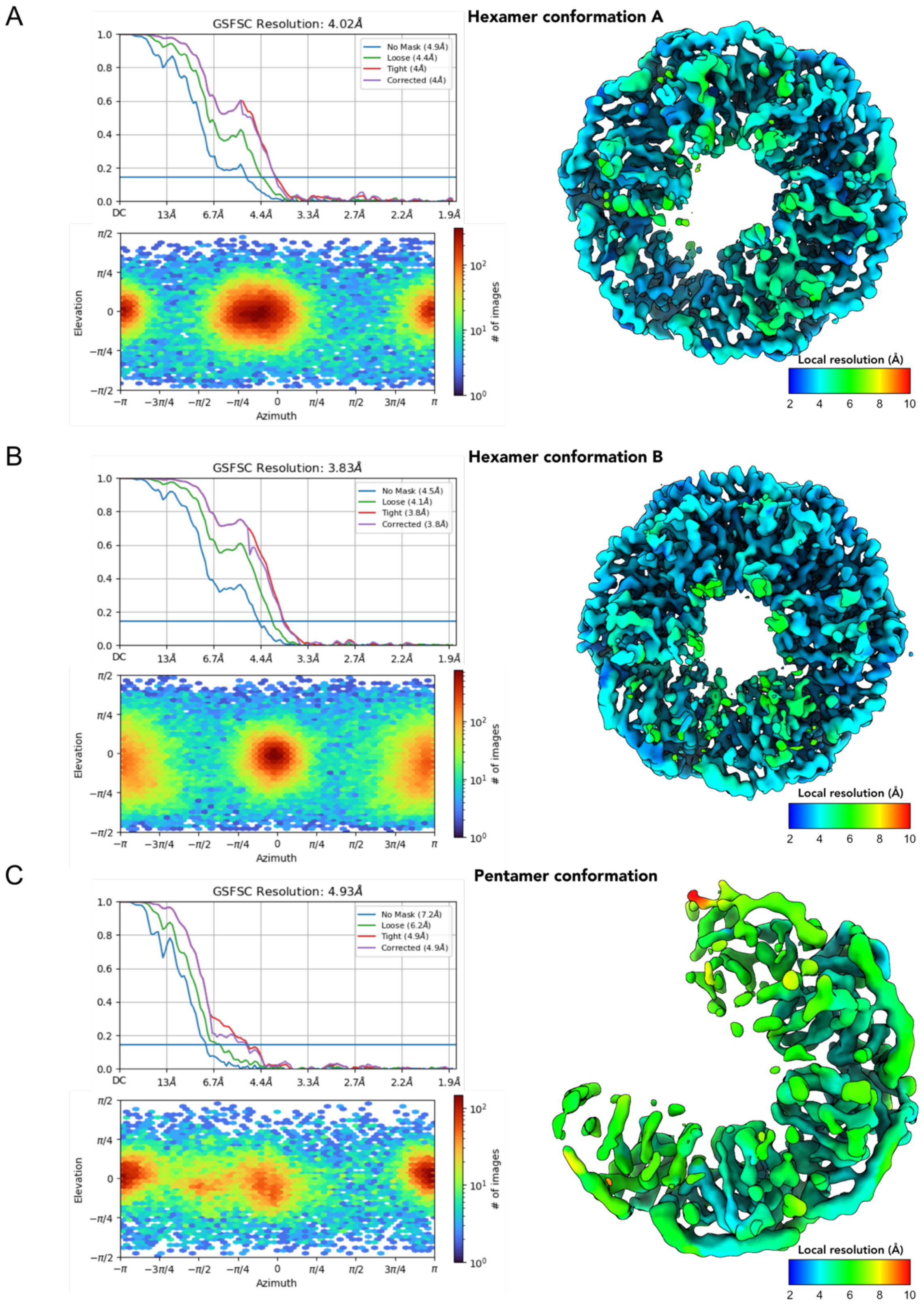

**Figure S2.** Validation of ChlI cryo-EM structures. A-C) Data for the hexamer conformation A (A), hexamer conformation B (B), and pentamer conformation (C). In each panel, the Gold Standard Fourier Shell Correlation (GSFSC) with the blue horizontal line marking the GSFSC value of 0.143 is shown in the top left graph, the angular distribution plot of the particles used for the reconstruction is shown in the bottom left graph, and the local resolution estimate of the ChlI cryo-EM map top view, calculated in CryoSPARC (see color code bar), is shown on the right.

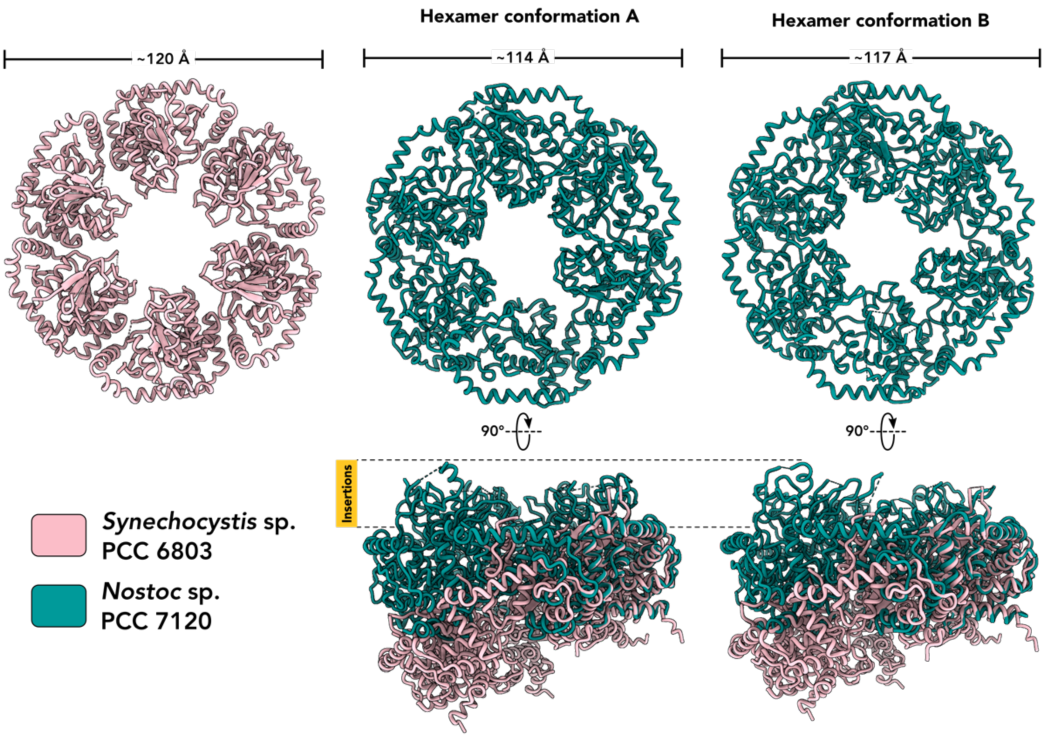

**Figure S3.** Comparison of cryo-EM hexamer structures of ChlI from *Nostoc* with the published X-ray structure of ChlI from *Synechocysts*. Top row: ribbon representations of the structure of *Synechocysts* ChlI (PDB: 6L8D, pink) and the *Nostoc* ChlI structures (this study, teal) viewed from above. Approximate ring dimensions are shown; Bottom row: superposition of the *Nostoc* ChlI structures with the *Synechocystis* ChlI structure using the "matchmaker" command in ChimeraX applied to individual monomers of each structure. The positions of the insertions (orange) in *Nostoc* ChlI are indicated in the bottom row.

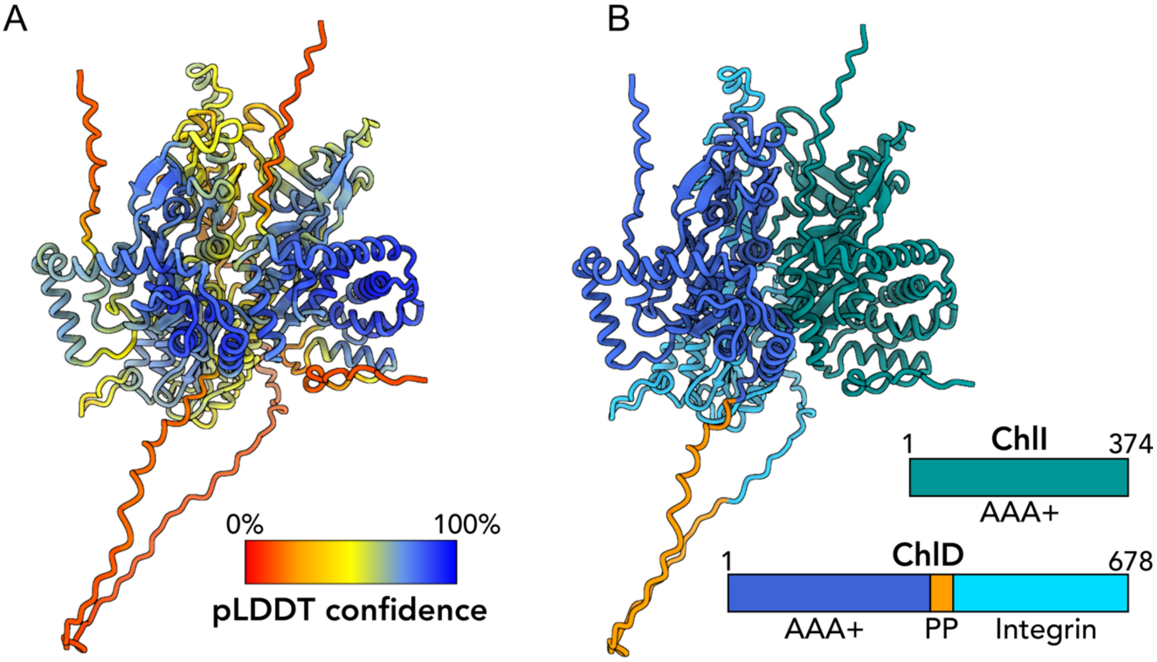

**Figure S4.** AlphaFold modeling of ChlI interactions with the partner protein ChlD. A) Ribbon representation of a ChlI-ChlD complex model colored by the pLDDT confidence of the structure prediction (see color code bar); B) Predicted complex of ChlI and ChlD viewed as in (A) and colored by protein domains. Schematic overview of ChlI and ChlD protein sequences is shown. The integrin domain (light blue) and the polyproline region (orange), which are characteristic for the ChlD protein, are indicated in the schemes and in the structure representation.
